# The fungal pathogen *Rhizoctonia solani* AG-8 has two nuclear haplotypes that differ in abundance

**DOI:** 10.1101/2025.06.30.662474

**Authors:** Jana Sperschneider, Kathleen DeBoer, Karam B Singh, Gupta V S R Vadakattu, Jonathan Anderson

**Author notes:** Department of Agriculture, Fisheries and Forestry, Canberra, ACT, Australia.

## Abstract

The fungal pathogen *Rhizoctonia solani* infects a diverse range of host plants and remains an intractable and economically significant disease for many crops. *R. solani* is classified into reproductively incompatible anastomosis groups (AGs). In the vegetative stage, most plant-pathogenic *R. solani* isolates are multinuclear and heterokaryotic, but little was previously known about the diversity between haplotypes due to highly fragmented, collapsed short-read assemblies. We present fully-phased, chromosome-scale genome assemblies of the broad host-range *R. solani* isolates AG8-1 and AG8-3. We demonstrate that both AG8 isolates have two distinct haplotypes, each of which is ∼50 Mbp spread across 16 chromosomes and use PacBio Iso-Seq data to achieve a high-quality gene annotation. We show that the two nuclear haplotypes display high heterozygosity and differences in haplotype abundance in vegetative cultures. Using transcriptome sequencing during infection of three types of host plants, we show that the less abundant haplotype in both AG8-1 and AG8-3 harbour more genes up-regulated during infection. Taken together, these findings address some of the observed phylogenetic heterogeneity of AG-8 isolates and provide a platform to further dissect the mechanisms enabling this globally significant agricultural pathogen to inflict losses to a range of crop hosts.

## Introduction

*Rhizoctonia solani*, a fungal species complex encompassing both pathogenic and non-pathogenic forms, is a significant threat to forestry and agriculture worldwide (Nagaraj et al. 2017). *R. solani* is responsible for a range of diseases, including root, crown, and stem rot, as well as damping-off and wilting, affecting a wide variety of economically important crops (Nizamani et al. 2025; Anderson 1982). *R. solani* is a fungal species complex comprising multiple genetic groups known as anastomosis groups (AGs), which are classified based on their ability to anastomose, or fuse, with hyphae of isolates belonging to the same group. Currently, 13 distinct AGs have been identified, along with a group of bridging isolates, AG-BI (Carling et al. 2002). These AGs differ in various aspects, including morphological traits and host range and can be distinguished at the genetic level, further supporting their classification as separate sub-species (Sneh et al. 1996). *R. solani* AG-8 is the causal agent of Rhizoctonia root rot and bare patch of wheat and barley. AG-8 isolates typically possess a broad host range, attacking both cereals and dicot crops including those commonly rotated with wheat or barley such as chickpea, lentil, lupin, and canola (Khangura et al. 1999; Sweetingham et al. 1986). Isolates of the AG-8 group result in losses of approximately $137 million annually in Australian cereal production (Murray and Brennan 2010; Huberli et al. 2025), over $100 million in Washington state, and billions of dollars globally (Okubara et al. 2014). While chemical control options exist in some crops, the availability of effective and reliable agrochemicals is limited thereby necessitating the adoption of alternative management strategies. Currently, crop protection largely relies on practices such as crop rotation, tillage, and, where possible, the use of resistant crop varieties (Okubara et al. 2014). These approaches are vital in mitigating the impact of *R. solani* and improving crop resilience and production against this widespread pathogen.

In natural environments, *R. solani* primarily reproduces asexually, existing mainly as vegetative mycelium or sclerotia, which serve as resting structures that enhance its survival and persistence in soil and plant debris (Anderson 1982). Isolates within the same AG may be able to form heterokaryons resulting from exchange of genetic material during anastomosis (Bolkan 1974). As a result, the AGs of *R. solani* can largely be considered reproductively isolated, each functioning as a distinct sub-species with specific ecological niches and pathogenic capabilities. Many of the highly pathogenic isolates of *R. solani* are multinuclear heterokaryotes, maintaining separate nuclei containing substantial diversity between the haplotypes. The heterokaryotic state creates difficulties in assembling genomes and assigning reads, particularly short reads, to any one individual haplotype. Thus, genomes assemblies from multinucleate isolates to date have mostly assembled haploid representations of the heterokaryotic state.

Currently, 24 genome assemblies exist for *R. solani* with isolates originating from AG1-1, AG2-2, AG-3, AG-4, AG-6, and AG-8 (Hane et al. 2014; Cubeta et al. 2014; Nadarajah et al. 2017; Wibberg et al. 2015, 2016, 2017; Ghosh et al. 2019; Zhang et al. 2021; Li et al. 2021; Kaushik et al. 2022; Lu et al. 2023; Liu et al. 2024; Xu et al. 2024). These draft genome assemblies exhibit a remarkable size range, with estimates from 33 Mbp for an AG1-IC isolate to approximately 70 Mbp for the AG3-1A1 and AG2-2IIIB consensus haploid assemblies (Wibberg et al. 2016; Kaushik et al. 2022). The diploid assembly of the uninucleate AG1-IA JN isolate was the largest to date at 97 Mbp (Li et al. 2021). This genomic variation underscores the complexity of *R. solani* at the molecular level and highlights the challenges inherent in studying its diverse genomic architecture. The genome data reported herein is the first chromosome-level assembly of haplotype genomes from multinuclear heterokaryotic *R. solani* isolates, which enables accurate analysis of intra-isolate diversity and the involvement of the homoeologous genes in the infection process across the broad range of hosts infected by AG-8.

## Methods and materials

### Growth of fungal isolates and inoculation of plant hosts

The *R. solani* isolate AG8-1 (WAC10335) was described previously (Hane et al. 2014). The AG8-3 isolate (WAC9760) was originally isolated from ‘bare patch’ affected wheat in Esperance Downs, Western Australia, (MacNish and Sweetingham, 1993) and was supplied by The Department of Primary Industries and Regional Development, Western Australia. Initial cultures were grown on potato dextrose agar (PDA) at 25ºC for five days. For inoculation of plant hosts a plug of agar from PDA cultures of AG8-1 and AG8-3 was used to inoculate sterile millet seed and incubated at 24ºC for 14 days. Millet cultures were dried in the laminar flow for 24 hours prior to freezing at -80ºC until use. Five millet seed were used to inoculate moist soil in 0.8 L pots and the pots incubated at 24ºC for 7 days, non-inoculated control pots were treated similarly without the addition of infected millet seeds. Four wheat (cv. Wyalkatchem), narrow leaf lupin (c.v. Coyote) or canola (cv. Zircon) seeds were planted into each pot to a depth of 25 mm and incubated at 16ºC with 12 hours of light per day at 350 lum. Plants were scored and photographed at 35 days after inoculation. For RNA-seq analysis of AG8-1 and AG8-3 infections of wheat, *Medicago truncatula, Brassica napus* and *Arabidopsis thaliana*, vermiculite was pre-infected with four Rhizoctonia-infected millet seed per pot for one week at 24ºC prior to planting. Wheat (cv Chinese Spring), *M. truncatula* (A17) and *Brassica napus* (cv Westar) seeds were surface sterilised with 70% ethanol, rinsed in sterile water and germinated on moist filter paper at 4ºC for 4 days prior to planting into *R. solani* pre-infected or non-infected (control) pots. *Arabidopsis thaliana* (Col-0) seeds were grown in vermiculite at 21ºC for 10 days prior to transplanting to *R. solani* pre-infected or non-infected (control) pots. All pots were then incubated at 16ºC for seven days prior to collection and washing of root tissue. Tissue was immediately frozen in liquid nitrogen and stored at -80ºC until RNA extraction.

### Isolation of genomic DNA, RNA and sequencing

A portion of hyphae from the growing tip of the colony was transferred to a liquid defined minimal medium (Solomon et al. 2004) with gentle shaking at 75 rpm at 22ºC for 10 days to enable growth of vegetative fungal hyphae in-vitro. Hyphal material was recovered from the culture by filtering through sterile cheesecloth and rinsing with sterile water prior to drying with sterile blotting paper. The hyphal material was frozen in liquid nitrogen and stored at -80ºC until DNA extraction. High molecular weight DNA was purified according to a protocol including the precipitation of small fragments using PEG (Debler et al. 2020). The purified DNA was subjected to PacBio HiFi sequencing by AGRF (Brisbane, Australia). For Hi-C sequencing, mycelium from liquid defined minimal medium cultures were cut into small fragments and cross-linked with 1% formaldehyde in PBS at room temperature for 20 minutes. Glycine was added to 125mM final concentration and incubated for 15 minutes prior to precipitation by centrifugation, washing in PBS and grinding in liquid nitrogen. Samples were sequenced by Phase Genomics (Seattle, USA). PacBio Iso-seq long read RNA sequencing data was obtained from vegetative mycelial cultures grown in potato dextrose broth (PDB) at 23ºC for 7 days with shaking at 75 rpm. RNA was extracted according to Anderson et al., 2017 and sequenced by AGRF (Brisbane, Australia). Stranded RNA-seq of vegetative mycelium and infected plant tissue was extracted (Anderson et al. 2017) and sequenced by Novogene (Hong Kong).

### Genome assembly and haplotype comparisons

The HiFi reads were downsampled with seqkit sample (--proportion 0.5) (Shen et al. 2016) and then assembled using hifiasm 0.19.6 in Hi-C integration mode and with default parameters (Cheng et al. 2021). Contaminants were identified using sequence similarity searches (BLAST 2.11.0 -db nt -evalue 1e-5 -perc_identity 75) (Altschul et al. 1990). HiFi reads were aligned to the assembly with minimap2 v2.22 (-ax map-hifi –secondary=no) (Li 2018) and contig coverage was called using bbmap’s pileup.sh tool on the minimap2 alignment file (http://sourceforge.net/projects/bbmap/). All contaminant contigs, contigs with less than 15x coverage and the mitochondrial contigs were removed from the assembly. Chromosomes were curated using visual inspection of Hi-C contact maps produced using Hi-C-Pro 3.1.0 (MAPQ 10) (Servant et al. 2015) and Hicexplorer 3.7.2 (Ramírez et al. 2018). The two haplotypes were compared to each other with mummer 4.0.0rc1, using nucmer and dnadiff (Marçais et al. 2018). Haplotype abundance was estimated by mapping the PacBio HiFi reads to the assemblies with minimap2 (-ax asm20 --secondary=no) and calculating coverage with bbmap’s pileup.sh (http://sourceforge.net/projects/bbmap/).

### Annotation and RNA-seq analysis

De novo repeats were predicted with RepeatModeler 2.0.2a and the option -LTRStruct (Flynn et al. 2020) and RepeatMasker 4.1.2p1 (-s -engine ncbi) (http://www.repeatmasker.org) was run with the RepeatModeler library to obtain statistics about repetitive element content. For gene annotation, RNA-seq reads were cleaned with fastp and default settings (Chen et al. 2018) and aligned with STAR 2.7.9a in 2-pass mode (-alignIntronMin 5 --alignIntronMax 3000 --alignMatesGapMax 3000 --outFilterMultimapNmax 100) which removes junctions supported by <= two reads (Dobin et al. 2013). Transcripts were assembled from the alignment with Stringtie 2.2.1 (-s 1 -m 200) with the appropriate strand setting (Pertea et al. 2015). Iso-Seq reads were cleaned with isoseq refine (https://github.com/PacificBiosciences/pbbioconda) and fastp (--trim_poly_x --adapter_fasta) (Chen et al. 2018). Clean Iso-Seq reads were aligned with minimap2 2.25 (-ax splice:hq -uf -G 3000) and transcripts were assembled with Stringtie (-L -m 200). The resulting sets of RNA-seq and Iso-Seq transcripts were merged into a consensus set with Stringtie (--merge). CodingQuarry 2.0 was run in pathogen mode with these transcripts (Testa et al. 2015). Funannotate 1.8.13 was run in training mode with the RNA-seq reads and Iso-Seq transcripts as input. We then ran funannotate predict (–ploidy 2 –optimize_augustus –busco_seed_species ustilago) and supplied the CodingQuarry predictions with the option -other_gff and set the weight to 1. BUSCO completeness was assessed on the annotated proteins using version 5.4.7. For gene expression analysis, we used Salmon 1.10.1 in genome decoy mode (Patro et al. 2017). We used Tximport (type = “salmon”) and DESeq2 to assess gene differential expression following default settings (*P*_adj□_< □0.1) (Love et al. 2014). Orthofinder 2.5.4 was used to extract 1-1 orthologs. We used DGenies and KaryoploteR for visualization of the chromosomes (Cabanettes and Klopp 2018; Gel and Serra 2017)

### Phylogenetic tree

We downloaded all publicly available *Rhizoctonia solani* genome assemblies from NCBI and used PHAME 1.0.5 (Shakya et al. 2020) to generate a maximum likelihood phylogeny which was visualized in iTOL (Letunic and Bork 2007).

## Results and Discussion

### Chromosome-scale, nuclear-phased assemblies for *Rhizoctonia solani* AG8-1 and AG8-3

*R. solani* AG8 has been reported to carry multiple nuclei per cell with thus far unknown ploidy. To resolve its haplotype genomes, we sequenced the broad host-range *R. solani* isolate AG8-1 (WAC10335) and another isolate, AG8-3. Both AG8-1 and AG8-3 cause bare patch and root-rot diseases on cereals and legumes with AG8-1 also impacting survival of canola seedlings (Sweetingham and MacNish 1991; Anderson et al. 2010; Foley et al. 2016; Kidd et al. 2021) (Supplementary Figure S1) For both isolates, we used PacBio HiFi and Hi-C data with hifiasm (Cheng et al. 2021) to assemble the genomes (AG8-1: 13.1 Gb HiFi reads and 16.5 Gb Hi-C reads; AG8-3: 12.2 Gb HiFi reads and 12.3 Gb Hi-C reads). This resulted in two haplotype assemblies of ∼50 Mbp each (Table 1), which we scaffolded into 16 chromosomes per haplotype. For AG8-3, two contigs displayed phase switches which we corrected prior to scaffolding. In the AG8-1 assembly, haplotype A contains only a single gap, whereas haplotype B was assembled completely from telomere to telomere (Table 1). In contrast, the AG8-3 assembly has five gaps in haplotype A and two in haplotype B. The unplaced contigs comprise only ∼1.7 Mbp per isolate and are lower in GC content. Hi-C contact maps of the curated chromosomes display a clear nuclear phasing signal (Figure 1A,B). The 16 chromosomes are highly collinear, both within each isolate and between the two isolates (Figure 1C,D,E). These assemblies are a vast improvement compared to the short-read assembly of isolate AG8-1, which was a highly fragmented, haploid representation of the two haplotypes (39.8 Mbp, 857 scaffolds with 13.1% gaps, L50:160.5 Kbp) (Hane et al. 2014).

**Table 1:** Genome assembly statistics for *R. solani* AG8-1 and AG8-3.

|  | AG8-1 |  |  | AG8-3 |  |  |
| --- | --- | --- | --- | --- | --- | --- |
|  | Haplotype A | Haplotype B | Unplaced contigs | Haplotype A | Haplotype B | Unplaced contigs |
| Assembly size | 51.025 Mbp | 49.182 Mbp | 1.688 Mbp | 50.580 Mbp | 49.486 Mbp | 1.653 Mbp |
| No. of scaffolds | 16 | 16 | 17 | 16 | 16 | 24 |
| Number of gaps | 1 | 0 | 0 | 5 | 2 | 0 |
| GC content | 48.45% | 48.47% | 43.83% | 48.43% | 48.44% | 41.78% |
| No. of telomeres (forward/reverse) | 12/14 | 15/14 | 2/0 | 13/13 | 15/14 | 1/2 |
| Annotated genes | 14,941 | 14,594 | 217 | 15,260 | 15,159 | 171 |
| Complete BUSCOs | 98.8% | 99.3% | 0.1% | 97.7% | 97.5% | 0.3% |
| Duplicated BUSCOs | 0.9% | 0.5% | 0.1% | 0.7% | 0.5% | 0% |
| Repeat content | 12.81% | 13.24% | 19.09% | 13.96% | 13.21% | 24.75% |
| Retroelements | 10.44% | 11.03% | 11.81% | 11.89% | 11.34% | 14.21% |
| DNA transposons | 1.80% | 1.63% | 4.06% | 1.42% | 1.32% | 2.04% |

**Figure 1:**
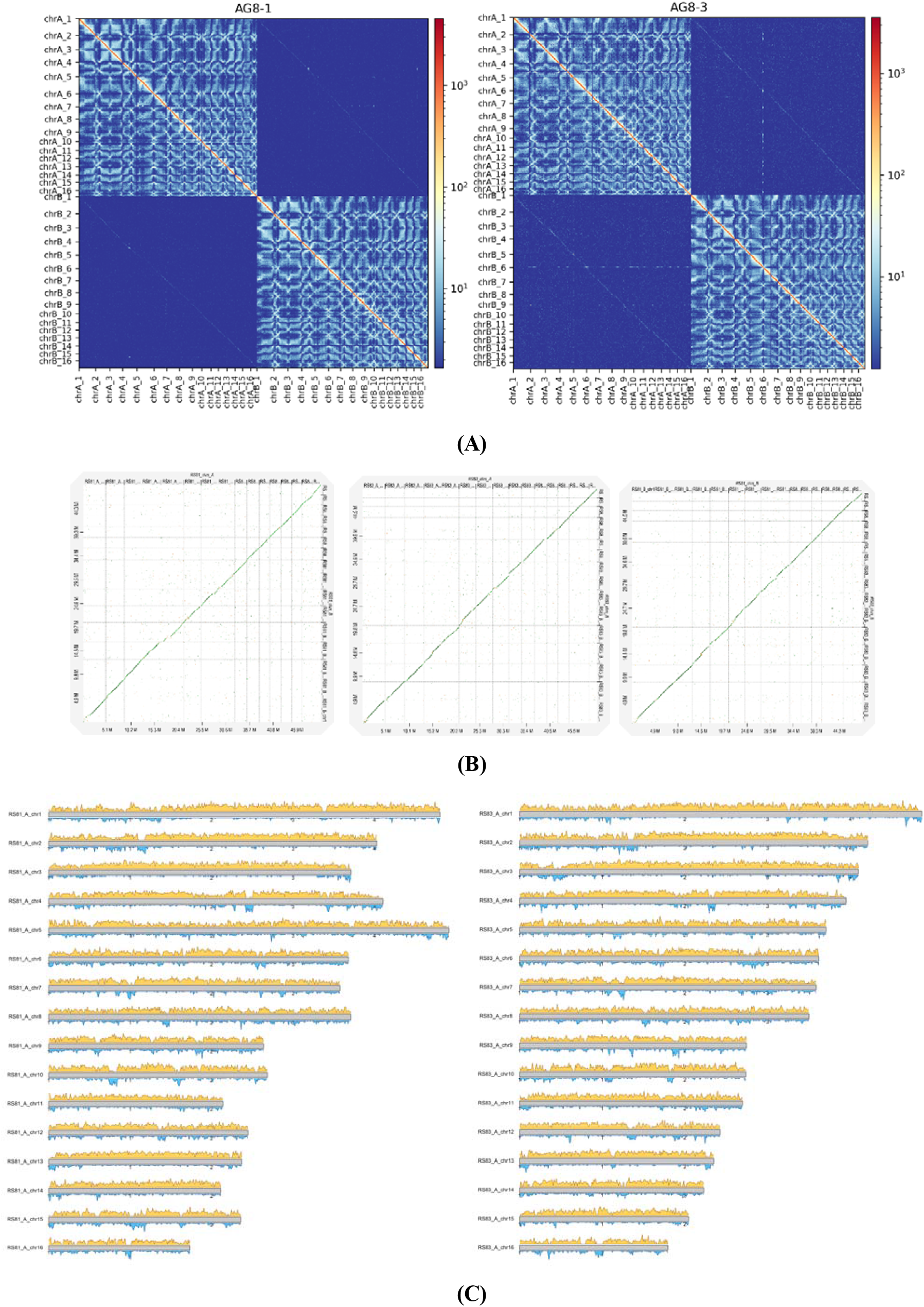
Hi-C contact maps and synteny of the 8-1 and 8-3 chromosomes. **(A)** Hi-C contact maps of isolates 8-1 and 8-3 show two distinct nuclear compartments. **(B)** Dot plots show that the chromosomes of the isolates are highly syntenic. **(C**) Gene density and repeat densities (10Kb bins) are shown in yellow and blue, respectively, for one haplotype of AG8-1 and AG8-3.

Next, we annotated repetitive elements for each genome. All four haplotypes had ∼13% repetitive bases, with retroelements the dominant class of transposable elements (Table 1). We observed highly repetitive regions on each chromosome, some of which might correspond to centromeric regions (Figure 1C). To enable a highly accurate gene annotation, we generated PacBio Iso-seq data for both isolates from vegetative cultures and supplemented this with RNA-seq data from infection (Supplementary Table 1). This resulted in ∼15k genes per haplotype with over 98% BUSCO completeness (Table 1).

**Supplementary Table 1:** Mapping rates for RNA-seq data sets.

| RNA-seq sample | Isolate | Number of reads | % mapped to AG8-1 | % mapped to AG8-3 |
| --- | --- | --- | --- | --- |
| Arabidopsis 7dpi Rep1 | AG8-1 | 96,471,543 | 5.35% | - |
| Arabidopsis 7dpi Rep2 |  | 94,072,660 | 2.74% | - |
| Arabidopsis 7dpi Rep3 |  | 96,721,875 | 8.84% | - |
| Canola 7dpi Rep1 |  | 41,278,768 | 3.31% | - |
| Canola 7dpi Rep2 |  | 32,206,582 | 4.31% | - |
| Canola 7dpi Rep3 |  | 38,579,162 | 2.96% | - |
| Medicago 7dpi Rep1 |  | 90,295,724 | 3.97% | - |
| Medicago 7dpi Rep2 |  | 96,589,990 | 3.24% | - |
| Medicago 7dpi Rep3 |  | 97,443,323 | 6.64% | - |
| Wheat 7dpi Rep1 |  | 35,228,760 | 9.81% | - |
| Wheat 7dpi Rep2 |  | 49,137,925 | 8.01% | - |
| Wheat 7dpi Rep3 |  | 36,486,393 | 18.95% | - |
| Wheat 7dpi Rep1 | AG8-3 | 31,965,766 | - | 6.71% |
| Wheat 7dpi Rep2 |  | 45,683,910 | - | 9.78% |
| Wheat 7dpi Rep3 |  | 47,054,443 | - | 13.57% |

### Both *Rhizoctonia solani* AG8-1 and AG8-3 have two nuclear haplotypes that are highly heterozygous

Next, we investigated the differences between the two nuclear haplotypes for each isolate (Table 2). The haplotypes of isolate AG8-1 are highly heterozygous with ∼9% unaligned bases, 96.8% average identity of alignments and 880K SNPs. The haplotypes of isolate AG8-3 are less heterozygous but still substantially different with ∼7% unaligned bases, 98% average identity of alignments and 551K SNPs. Interestingly, the AG8-1 haplotype A is substantially different to all other three haplotypes whereas the AG8-1 haplotype B is more similar to the AG8-3 haplotypes than to the AG8-1 haplotype A (Table 2). A phylogenetic tree of publicly available *Rhizoctonia* genome assemblies confirms that the AG8-1 haplotype B is more closely related to the AG8-3 haplotypes than to the AG8-1 haplotype B (Figure 2). Furthermore, the placing of the short-read assembly of AG8-1 in the phylogenetic tree confirms that it is a collapsed representation of the two haplotypes (Hane et al. 2014). We further investigated the SNP content between the two haplotypes to assess if Repeat-Induced Point (RIP) mutations are present as indicated previously (Hane et al. 2014). Using the RIPper software (van Wyk et al. 2019), only 0.07% and 0.09% of the AG8-1 and AG8-3 genomes were estimated to be affected by RIP, respectively. However, it may be possible that a RIP-like process might operate in *Rhizoctonia* that is different from the one in ascomycetes.

**Table 2:** Haplotype alignment and SNP statistics.

| SNPs \| Alignment identity | AG8-1A | AG8-1B | AG8-3A | AG8-3B |
| --- | --- | --- | --- | --- |
| AG8-1A | - | 880,333 \| 96.82% | 890,832 \| 96.77% | 902,196 \| 96.74% |
| AG8-1B |  | - | 551,518 \| 98.11% | 557,047 \| 98.09% |
| AG8-3A |  |  | - | 551,138 \| 98.11% |
| AG8-3B |  |  |  | - |

**Figure 2:**
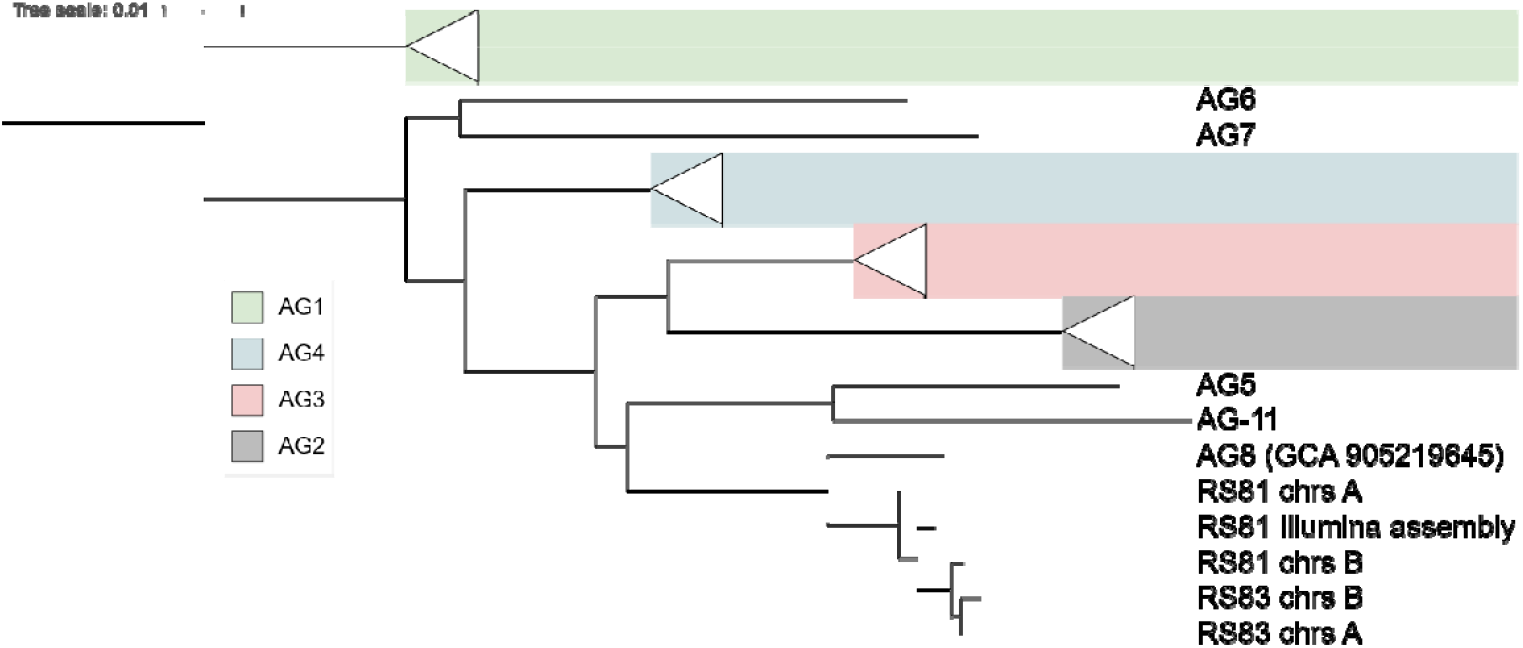
A phylogenetic tree of Rhizoctonia isolate genomes. Rhizoctonia isolates group according to their AG groups.

**Supplementary Figure S1.**
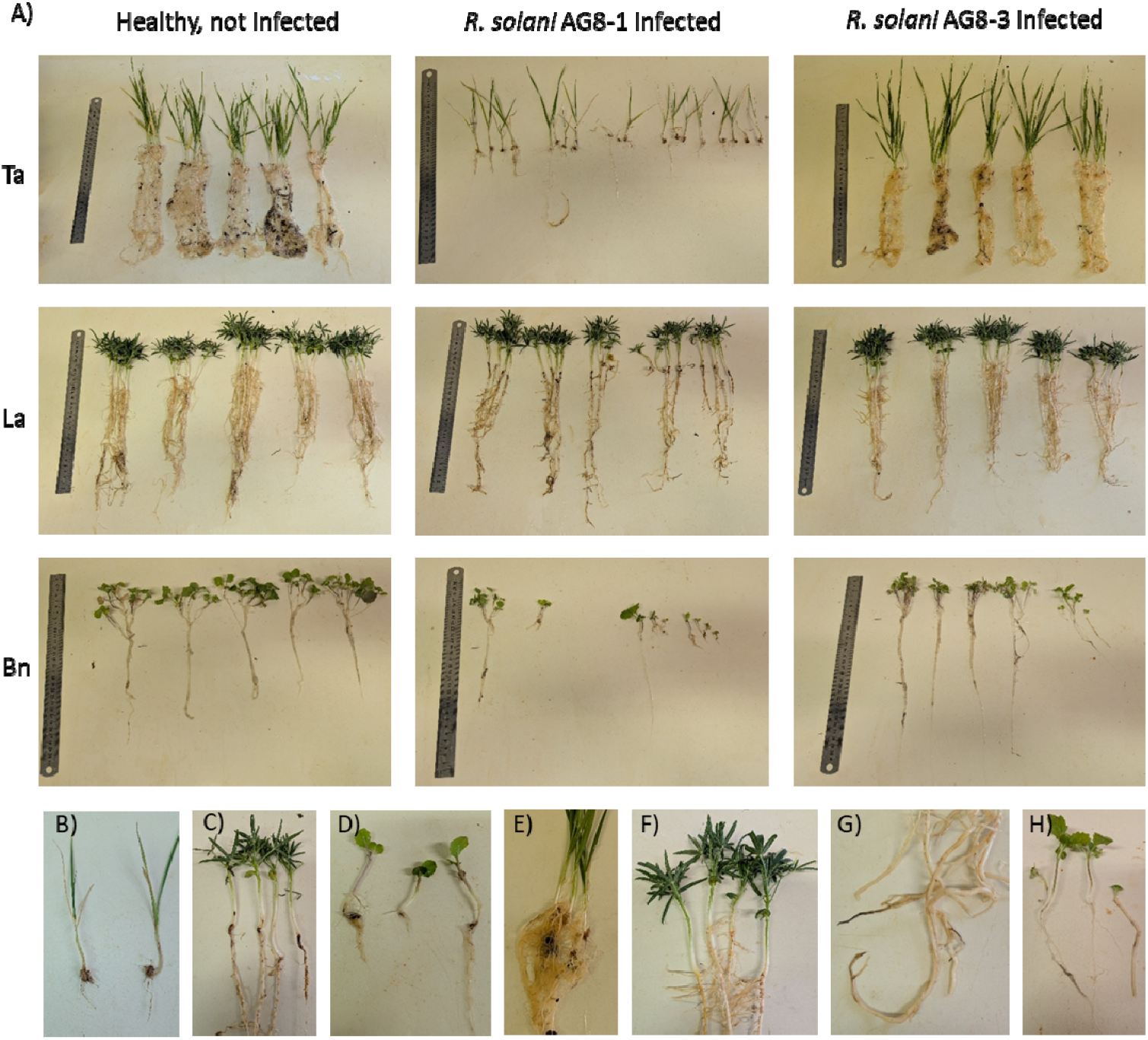
Symptoms of infection with *R. solani* AG8-1 and AG8-3. A) Healthy, AG8-1 or AG8-3 infected wheat (*Triticum aestivum*, Ta), narrow leaf lupin (*Lupinus angustifolius*, La) and canola (*Brassica napus*, Bn). Symptoms of infection with AG8-1 on wheat (B), narrow leaf lupin (C) and canola (D). Symptoms of infection with AG8-3 on wheat (E), narrow leaf lupin (F, G) and canola (H).

### Haplotype abundance and transcriptome contribution of co-existing genomes during plant infection

Mapping of the PacBio HiFi reads back to the assemblies indicated that the two haplotypes occur at significantly different abundances in vegetative cultures (Figure 3). For AG8-1, the A haplotype is more abundant than the B haplotype (8.5% higher). In contrast, for AG8-3 the B haplotype is more abundant than the A haplotype (9.9% higher). To compare gene expression between the two haplotypes, we conduced a protein orthology analysis. We selected single-copy orthologues that differentiated by at least one SNP in their coding sequences to assess gene expression on the two haplotypes (AG8-1: *n =* 10,514; AG8-3: *n =* 9,873). We used RNA-seq data from infection of *Triticum aestivum* with AG8-1 and AG8-3 as well as infection of *Arabidopsis thaliana, Medicago truncatula* and *Brassica napus* with AG8-1 (Supplementary Table 1). Principal component analysis (PCA) of RNA-seq data shows that samples all clustered according to biological conditions (Supplementary Figure 2).

**Figure 3:**
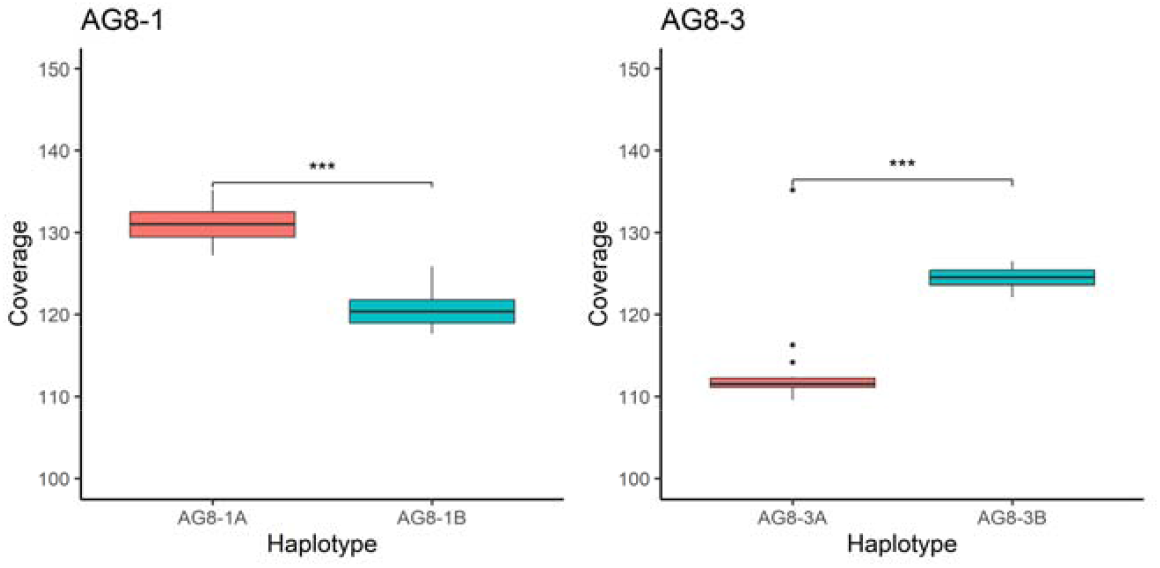
Haplotype abundance indicated by chromosome read coverage for the two isolates.

**Supplementary Figure 2:**
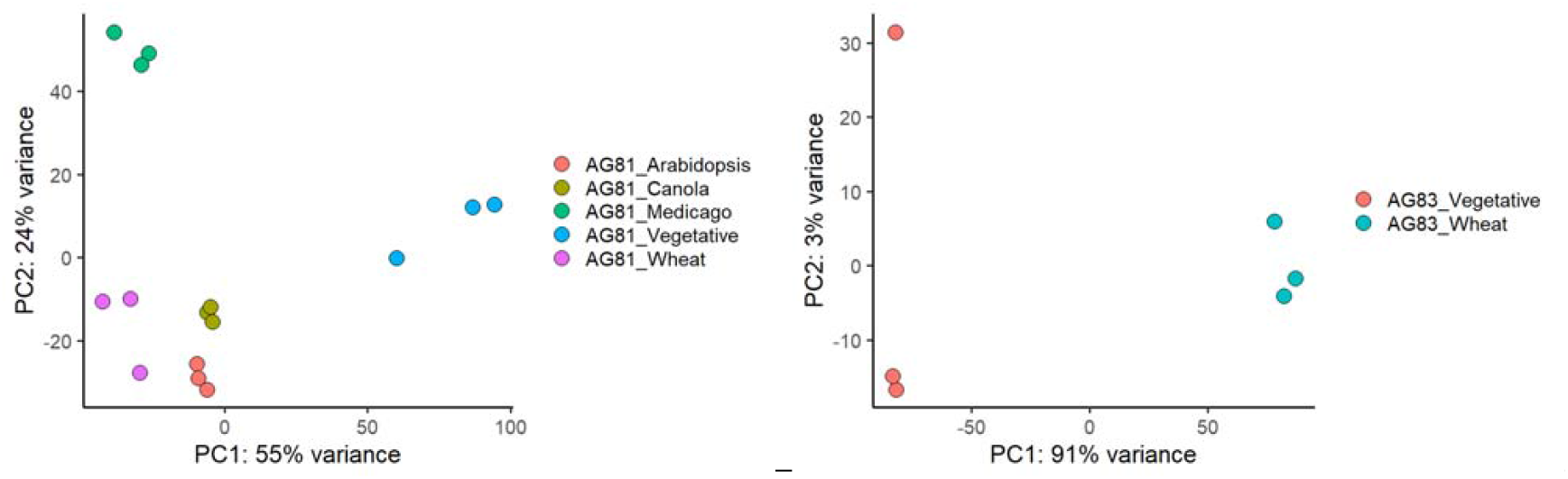
PCA plots for the RNA-seq data sets.

When comparing transcription between haplotypes, for AG8-1 the more abundant A haplotype has significantly higher transcription than the B haplotype for all conditions except during wheat, canola (*Brassica napus*) and barrel medic/medicago (*Medicago truncatula*) infection (Figure 4). Similarly, the more abundant B haplotype has higher transcription than the A haplotype for AG8-3 (Figure 4). However, gene regulation does not follow the same trend of haplotype abundance. In AG8-3, 67.1% of genes up-regulated during infection of wheat are located on the less abundant A haplotype (Figure 4) suggesting a potential for greater involvement of the genes contained within this haplotype to be important for infection of wheat. The AG8-1 isolate showed a lower degree of variance in haplotype expression with only a slight over-representation of the less abundant B genotype during infection of wheat, medicago and canola (Figure 4). Homeolog expression dominance has previously been observed between the sub-genomes of the uninucleate AG1-IA JN strain (Li et al. 2021). Further analysis of the differentially expressed (DE) homeolog pairs in the JN strain suggested less evolutionary constraint of the DE genes than homeolog pairs with similar expression profiles (Li et al. 2021). This suggested that gene duplication can lead to genetic divergence, potentially acting as a source of variation in pathogenicity and may contribute to the unusually broad host range of AG-8 isolates.

**Figure 4:**
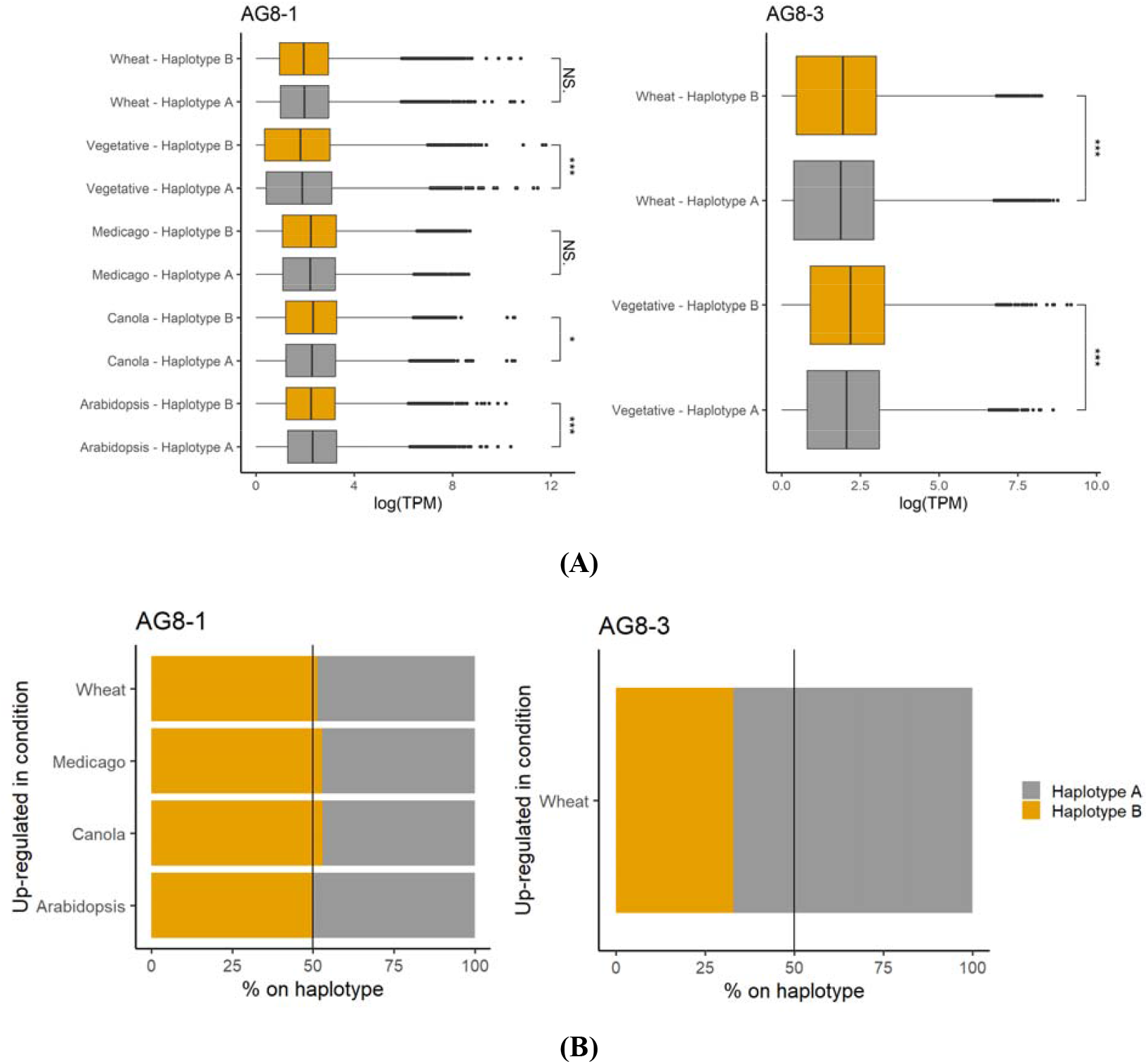
RNA-seq expression analysis during vegetative stages and host infection. **(A)** Significantly higher TPM values are recorded for the more abundant haplotype for AG8-1 (haplotype A) and AG8-3 (haplotype B) in most conditions. **(B)** In contrast, a higher proportion of genes up-regulated during plant infection reside on the A haplotype with lower abundance in AG8-3. The AG8-1 isolate showed a slight over-representation of the less abundant B haplotype during infection of wheat, medicago and canola.

Relatively little is known about the genomic characteristics and pathogenicity genes of disease-causing isolates of *R. solani*. Anderson et al., (2017) compared the secretomes of AG1-1, AG3 and AG-8 isolates and predicted conserved and unique putative secreted effector proteins including a large array of cell wall degrading enzymes (CAZYs) but a relatively small number of putative secreted effectors compared to other plant pathogenic fungal species. However, as these genome assemblies were haploid representations of heterokaryotic isolates, the genetic diversity hidden within the homeologs could not be investigated. Similarly, Ghosh et al., (2019) sequenced the genomes of two AG1-IA strains highly virulent for causing rice sheath blight in India and compared these to the pre-existing AG1-IA genome. The highly virulent isolates showed expansion or emergence of orthogroups including pectate lyases, glycoside hydrolases, UDP-glucuronosyltransferases, oxidation-reduction-related genes such as NADPH-dependent oxidoreductase, cytochrome P450 and zinc finger transcription factors. Further comparative genomics reported by Liu et al., (2024) identified a protein domain conserved among *R. solani* genome sequences from diverse anastomosis groups and only present in basidiomycete species. Silencing of the encoding genes in the AG1-ZJ isolate reduced lesion size on maize and topical RNAi treatment targeting the *R. solani* genes reduced lesion size on maize, rice and wheat, suggesting the domain may be important for pathogenicity across AGs and potentially on diverse hosts.

## Conclusion

The generation of high-quality chromosome level genome assemblies that differentiate haplotypes and enable exploration of the role of these during infection of diverse host crops is an important step to improve crop protection. As a heterokaryotic, multinuclear fungus little was previously known about the diversity between haplotypes in *R. solani* AG-8. Here we present chromosome-level, nuclear-phased assemblies for two *R. solani* AG-8 isolates from Australian grain production areas, which are pathogenic on cereals, brassicas and legumes. This information gives a good foundation for future research to understand the ecology and epidemiology of *R. solani* AG8 across diverse host genotypes and environments. We anticipate that this new resource will facilitate future discoveries in this pervasive and important fungal pathogen.

## Data availability statement

The AG8-1 vegetative and wheat infection RNA-seq reads are available under NCBI BioProject PRJNA371695. The AG8-1 Medicago infection RNA-seq reads are available under NCBI PRJNA369210 and SRP098557. All other data including the genome assemblies, annotations, PacBio Iso-Seq data and other RNA-seq data is available at the CSIRO Data Access Portal https://data.csiro.au/collection/csiro:65845.

## Acknowledgements

The authors acknowledge Scott Rice and the CSIRO Microbiomes for One System Health Future Science Platform for providing funding to support genome sequencing activities, and Rhonda Foley, Gagan Garg and Nick Pain for advice and/or assistance with transcriptomic studies.

